# A Primary Neural Cell Culture Model for Neuroinflammation

**DOI:** 10.1101/2020.01.23.914226

**Authors:** Noah Goshi, Rhianna K. Morgan, Pamela J. Lein, Erkin Seker

## Abstract

Interactions between neurons, astrocytes and microglia critically influence neuroinflammatory responses to insult in the central nervous system. Studying neuroinflammation *in vitro* has been difficult because most primary culture models do not include all three critical cell types. We describe an *in vitro* model of neuroinflammation comprised of neurons, astrocytes and microglia. Primary rat cortical cells were cultured in a serum-free medium used to co-culture neurons and astrocytes that is supplemented with three factors (IL-34, TGF-β and cholesterol) used to support isolated microglia. This “tri-culture” can be maintained for at least 14 days *in vitro* while retaining a physiologically-relevant representation of all three cell types. Additionally, we demonstrate that the tri-culture system responds to lipopolysaccharide, mechanical trauma and excitotoxicity with both neurotoxic and neuroprotective aspects of the neuroinflammatory response observed *in vivo*. We expect the tri-culture model will enable mechanistic studies of neuroinflammation *in vitro* with enhanced physiological relevance.

## Introduction

Neuroinflammation is present in most, if not all, pathological conditions in the central nervous system (CNS), either acting as the primary driver of these conditions, or as a response to neurodegeneration or disruption of homeostasis following disease progression (Corps et al., 2015; Jayaraj et al., 2019; Waisman et al., 2015). Following insult or injury to the CNS, the two primary cell types associated with neuroinflammation, microglia and astrocytes (Eggen et al., 2013; Jensen et al., 2013), become activated, as indicated by changes in their morphology and phenotype (Polikov et al., 2005). Upon activation, astrocytes can undergo a transformation to a diverse range of phenotypes including highly neurotoxic “A1” phenotype or a neuroprotective “A2” phenotype depending on the mode of activation (Liddelow et al., 2017; Miller, 2018). Microglia were also thought to undergo a similar transformation into neurotoxic/neuroprotective “M1/M2” phenotypes; however, recent evidence suggests that microglial polarization is much more complex, with microglia displaying unique gene expression profiles depending on the activating stimuli (Li and Barres, 2017; Salter and Stevens, 2017). It is now accepted that neuroinflammation is a highly complex process that can be neurotoxic or neuroprotective depending on the disease state and progression. Therefore, there is significant interest in gaining a better understanding of the cellular and molecular cues that influence the phenotypic state of astrocytes and microglia.

While there are a number of *in vivo* models to study neuroinflammation, *in vitro* models are often used to investigate specific molecular pathways. *In vitro* models have been useful in gaining a better understanding of how microglia and astrocytes become activated towards neuroprotective or neurotoxic roles and of their impact on neuronal populations. In this paper, we report on an enhanced cell culture model comprised of all three major cell types associated with neuroinflammation – neurons, astrocytes and microglia. Current *in vitro* models of neuroinflammation typically consist of cultures of individual cell types with conditioned media from one cell type transferred to cultures of another cell type (Guttenplan and Liddelow, 2018; Harry and Kraft, 2008; Luna-Medina et al., 2005). While these models have provided insight into neuroinflammatory processes (Liddelow et al., 2017), these models contain inherent limitations, most notably the inability to observe the effect of membrane-bound or cell proximity-dependent mechanisms and the fact that the concentration of secreted cytokines transferred between the cultures may not be physiologically relevant. An alternative model involves seeding microglia over a previously established primary neuron culture to observe the effect of this cell-cell interaction over a short period of time (24-72h) (Culbert et al., 2006; Gresa-Arribas et al., 2012; Roque and Costa, 2017). In addition to the limited time-scale of this model, the culture media used to support the microglia prior to their addition to the neuronal culture contains a high concentration of serum, likely causing the microglia to be in an already activated state before their addition to the neurons (Bohlen et al., 2017).

To address the shortcomings of existing *in vitro* models of neuroinflammation, we developed a tri-culture model in which primary rat cortical cells were maintained in culture medium developed to support neuron-astrocyte co-culture media (Chapman et al., 2015) supplemented with interleukin-34 (IL-34), transforming growth factor beta (TGF-β) and cholesterol. These three factors have been previously shown to support isolated microglia cultures (Bohlen et al., 2017). We demonstrate that the tri-culture can be maintained for at least 14 days *in vitro* (DIV), without any deleterious effect of the continuous presence of microglia on the overall health of the neurons in the tri-culture. The tri-culture contains a similar relative percentage of neurons and displays a similar amount of neurite growth as compared to the microglia-free, neuron-astrocyte co-cultures. Furthermore, we demonstrate that the tri-culture system responds to several pro-inflammatory stimuli, including lipopolysaccharide (LPS), mechanical trauma, and excitotoxicity, in a manner similar to that observed *in vivo*.

## Results and Discussion

### The tri-culture media is able to support neurons, astrocytes and microglia *in vitro*

Primary cortical cells taken from neonatal rats were cultured in our previously described neuron-astrocyte co-culture media (Chapman et al., 2015), or in our tri-culture media consisting of the co-culture media supplemented with 100 ng/mL IL-34, 2 ng/mL TGF-β and 1.5 μg/mL cholesterol. Immunostaining for Iba1 revealed that there was a significant population of microglia at both DIV 7 and 14 in the cultures maintained in the tri-culture media, which are absent from cultures maintained in the co-culture medium (Figure 1A and B). A two-way ANOVA did not establish a significant interaction between the time in culture and media type on the number of microglia present in the culture (p = 0.44). Analysis of the main effects indicates that the tri-cultures have significantly more microglia present than the co-cultures (p = 0.0025); however, the time in culture did not impact the number of microglia (p = 0.44). The total number of microglia in the tri-culture accounts for ~7-8% of the total cell population in agreement with microglia numbers reported *in vivo* (Savchenko et al., 2000; von Bartheld et al., 2016). These microglia exhibit an amoeboid morphology and not the highly ramified morphology found by previous researchers culturing isolated microglia with these three supplements (Bohlen et al., 2017). We propose that this difference in morphology is due to the fact that the cortical cells were taken from neonatal rats, and at this developmental stage, microglia predominantly exhibit an amoeboid or primitive ramified morphology even under normal physiological conditions (Cuadros and Navascue, 1998; Dalmau et al., 2003; Harry and Kraft, 2012). This hypothesis is supported by the presence of galactin-3 in the tri-culture (Figure 5D). Galactin-3 has been shown to induce an amoeboid morphology in microglia, and the transcription factor for galactin-3 is highly expressed in healthy neonate amoeboid microglia, but not in adult microglia showing ramified morphologies (Reichert and Rotshenker, 2019).

**Figure 1:**
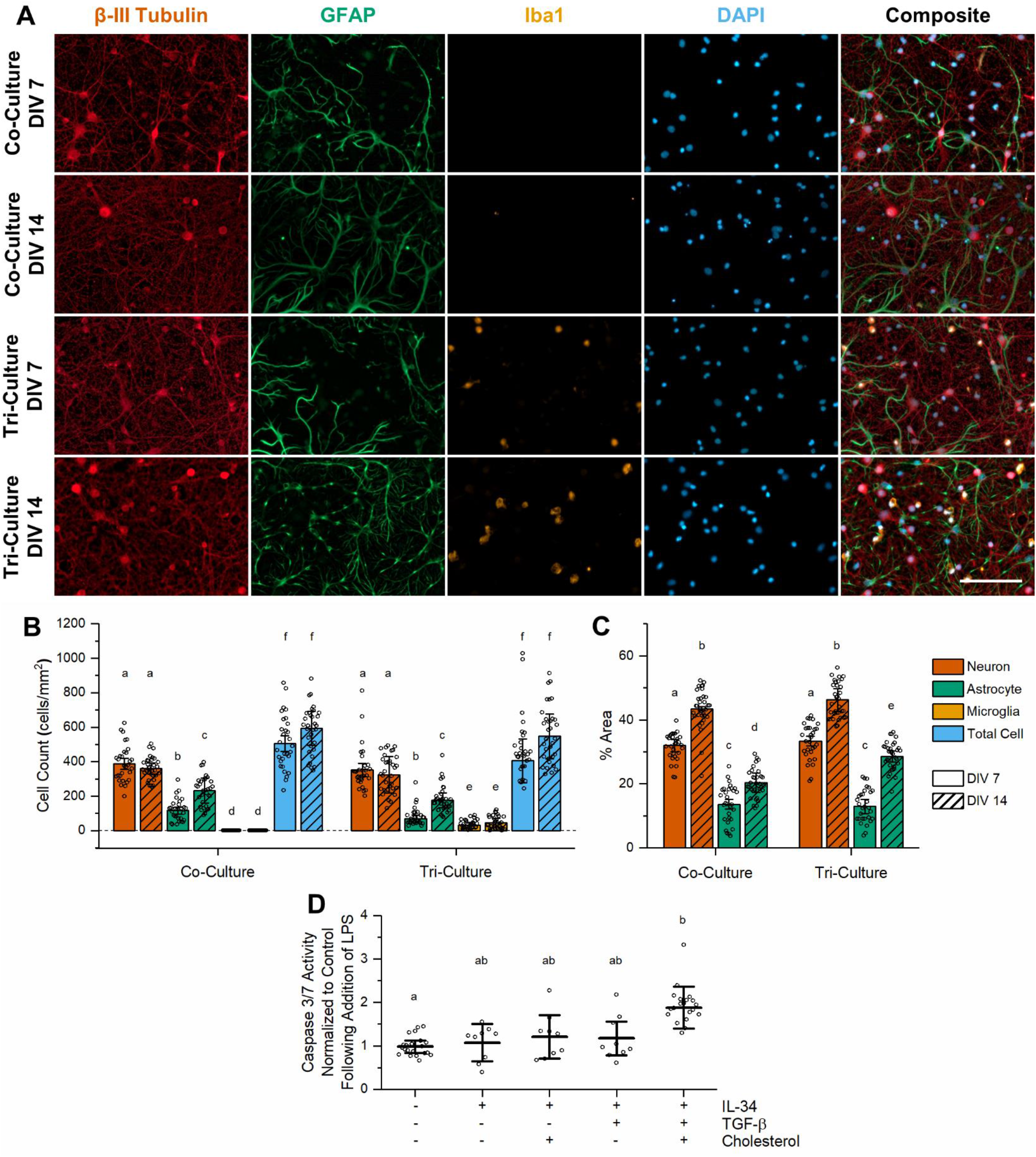
Tri-culture media is capable of supporting neurons, astrocytes and microglia. (**A**) Representative fluorescence images of the tri- and co-cultures at DIV 7 and 14. The cultures were immunostained for the three cell types of interest: neurons–anti-βIII-tubulin (red), astrocytes–anti-GFAP (green), microglia–anti-Iba1 (orange), general nuclear stain DAPI (blue). Microglia are present in the tri-culture at both time points, but are absent in the co-culture. Scale bar = 100 μm. (**B**) Quantification of the number of each cell type and total number of cells per mm2 for each condition shown in Figure 1A (mean ± SD, n=3). (**C**) Quantification of the percent area coverage of the neurons and astrocytes in the co- and tri-cultures at DIV 7 and 14 (mean ± SD, n=3). (**D**) Comparing the effects on apoptosis following a 48 h incubation with 5 μM LPS on DIV 7 cortical cultures maintained in different media types (mean ± SD, n=3-6). (**B-D**) The letters above the bars indicate statistically distinct groups (p < 0.05), while the points indicate the values of the technical replicates.

Immunostaining for β-III tubulin to label neurons and GFAP to label astrocytes revealed a healthy population of both neurons and astrocytes in both the tri- and co-cultures (Figure 1A). The number of neurons present was not affected by either the media type or time in culture (Figure 1B, p = 0.44 and p = 0.31 respectively). Additionally, neurite outgrowth (measured as the percent of the area stained for β-III tubulin as compared to the total image area) was not statistically different between the co- and tri-cultures (Figures 1C, p = 0.13), and the neurons continue to produce new projections through DIV 14 as determined by the significant increase in neuron percent area coverage from DIV 7 to 14 (p = 1.04 x 10^−5^). As no mitotic inhibitors were added to the culture media, the number of astrocytes significantly increased in both culture types from DIV 7 to 14 (Figure 1B, p = 0.0023). While the analysis of the main effects from the two-way ANOVA did not reveal a significant difference in astrocyte population between the two culture types (p = 0.062), the results suggest that the tri-cultures may contain a lower number of astrocytes than the co-cultures. This may be due to the presence of TGF-β in the tri-culture media, which has been shown to reduce astrocyte proliferation (Lindholm et al., 1992). Unlike the neurons, the astrocyte percent area coverage showed a significant interaction between the culture type and time in culture (Figure 1C, p = 0.0077), with the tri-culture having a similar astrocyte percent area coverage to the co-culture at DIV 7 (p = 0.90), but a significantly higher astrocyte percent area coverage at DIV 14 (p = 0.0076). This may be due in part to the large number of fine astrocyte processes seen in the DIV 14 tri-culture, which is overvalued by the auto-thresholding process used to calculate these values.

As the three media supplements present in the tri-culture were initially used to support isolated microglia, and were all derived from astrocyte conditioned media (Bohlen et al., 2017), we were interested to see if the neurons or astrocytes might be constitutively secreting some of these factors in the tri-culture, and thus making their addition to the tri-culture media redundant. An exploratory study unambiguously indicated that IL-34 was required for microglia survival (Figure 1 – figure supplement 1); however, the additional presence of TGF-β or cholesterol did not appear to impact the number of microglia present in the culture. Additionally, neither TGF-β nor cholesterol allowed for microglia survival on their own. Previous studies have indicated that the activation of the colony stimulating factor 1 receptor (CSF1R) via colony stimulating factor 1 (CSF1) or IL-34 is required for microglia viability both *in vitro* and *in vivo* (Chitu et al., 2016; Elmore et al., 2014; Erblich et al., 2011), and therefore its requirement in the tri-culture media is not unexpected.

Although the addition of TGF-β or cholesterol did not seem to be required for microglia viability in the tri-culture, it has been shown that TGF-β can induce a quiescent microglia phenotype *in vitro* (Abutbul et al., 2012), and it may even be required for cultured microglia to maintain their gene expression profile (Butovsky et al., 2014). Excess cholesterol in the culture media may also be beneficial for maintaining a functional microglia gene expression profile as it reduces the expression of apolipoprotein E (ApoE) (Mahley, 2016), which is critical for lipid transport, but it is also inversely correlated with the expression of microglia signature genes (Butovsky et al., 2014). Therefore, in order to determine if either TGF-β or cholesterol was required to maintain physiologically active microglia, we challenged cultures maintained under different combinations of TGF-β and cholesterol plus IL-34 with 5 μM of LPS. LPS is a potent activator of neuroinflammation and neuronal apoptosis, which acts through the toll-like receptor 4 (TLR4) found only on microglia in the CNS (Lehnardt et al., 2002; Nakamura et al., 1999). As expected, LPS did not increase caspase 3/7 activity in neuron-astrocyte co-cultures relative to the vehicle control. However, cultures grown in the presence of different combinations of the tri-culture factors responded to LPS with increased caspase 3/7 activity (Figure 1D, p = 0.011). In particular, cultures exposed to all 3 co-factors (IL-34, cholesterol and TGF-β) showed a significant increase in caspase 3/7 activity following the addition of LPS (p = 0.0081). Cultures maintained in the co-culture medium spiked with a subset of the co-factors (IL-34 alone, IL-34 plus cholesterol or IL-34 plus TGF-β) all showed increased caspase 3/7 activity, but a *post hoc* Tukey test did not reveal any significant differences between these cultures and either the co- or tri-culture. However, the fact that the change in caspase 3/7 activity of these cultures more closely resembled that of the co-culture (p = 0.99, 0.91 & 0.95 respectively) than that of the tri-culture (p = 0.064, 0.16, and 0.16 respectively), indicates that the microglia present in these cultures are most likely not physiologically active. Thus, all three factors are required to support a healthy tri-culture of neurons, astrocytes and microglia.

### Tri-culture response to LPS

The use of LPS to stimulate a neuroinflammatory response is used by researchers to study a wide range of neuroinflammatory and neurodegenerative conditions (Batista et al., 2019), including Alzheimer’s disease (AD) (Nazem et al., 2015), Parkinson’s disease (PD) (Dutta et al., 2008) and even mood disorders such as clinical depression (Henry et al., 2008). Therefore, we next determined the response of the tri-culture model to LPS. As stated previously, a 48 h incubation with 5 μM LPS significantly increased caspase 3/7 activity in the tri-culture model relative to the neuron-astrocyte co-culture (Figure 1D). This is in agreement with previous studies that show LPS induces a neurotoxic pro-inflammatory condition both *in vitro* and *in vivo* via apoptotic caspase-3 mediated mechanisms (Liddelow et al., 2017; Nimmervoll et al., 2013; Wang et al., 2005). However, while we do see some evidence of neurodegeneration and neurite loss in the LPS-exposed tri-cultures (Figure 2A), the extent of the damage is significantly less severe than what is observed *in vivo*. This may be because microglia require co-activation with both LPS and INF-γ (secreted by circulating leukocytes) to produce the severe neurotoxic effects seen *in vivo* (Papageorgiou et al., 2016). In the absence of leukocytes, no INF-γ is produced by the tri-culture in response to LPS (Figure 5 – figure supplement 1) leading to a less severe response.

**Figure 2:**
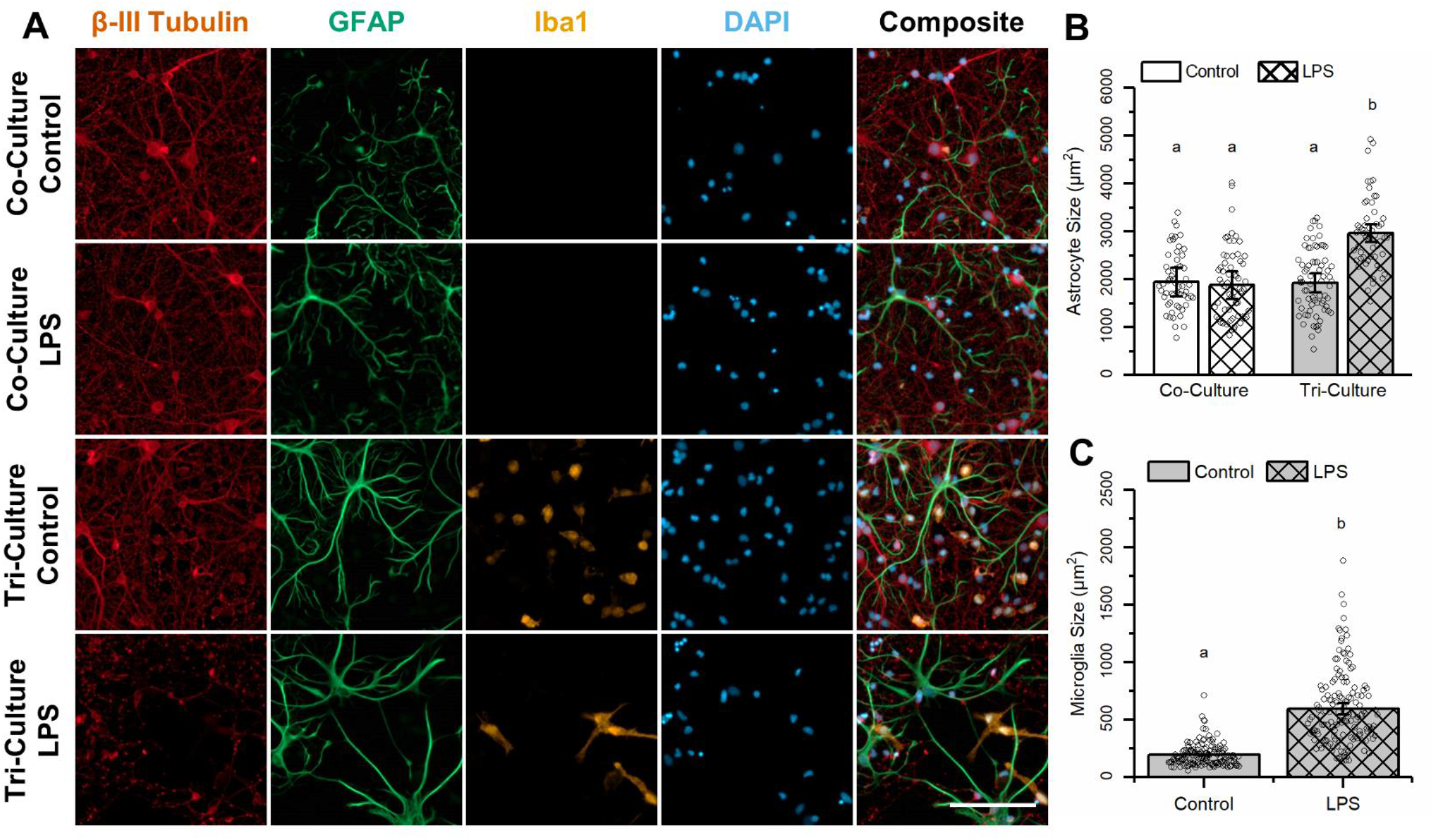
Comparing the effect of LPS on the tri- and co-culture models. Comparison of the effect of a 48 h exposure to 5 μM LPS on the tri- and co-cultures. (**A**) Representative fluorescence images of the tri- and co-cultures after a 48 h incubation with 5 μM LPS or vehicle. The cultures were immunostained for the three cell types of interest: neurons–anti-βIII-tubulin (red), astrocytes–anti-GFAP (green), microglia–anti-Iba1 (orange) and the general nuclear stain DAPI (blue). Scale bar = 100 μM. (**B**) A comparison of the average astrocyte area from the different conditions (mean ± SD, n=3). (**C**) Comparing the average microglia size in tri-cultures exposed to LPS vs. vehicle (mean ± SD, n=3). (**B-C**) The letters above the bars indicate statistically distinct groups (p < 0.05), while the points indicate the values of the technical replicates.

Immunohistochemical analyses (Figure 2A) reveal clear morphological changes in both astrocytes and microglia in the tri-culture after exposure to LPS. Astrocytes in the tri-culture showed reduced ramification, increased process length and hypertrophy, all of which are hallmarks of reactive astrocyte morphology (Schiweck et al., 2018). The increased process length and hypertrophy result in an overall increase in astrocyte area (Figure 2B) that was significantly higher in tri-cultures exposed to LPS relative to tri-cultures exposed to vehicle (p = 0.0030) or astrocytes in the co-culture exposed to LPS (p = 0.0039). Additionally, there was no change in the morphology or average area of astrocytes in the co-cultures exposed to LPS or vehicle (p = 0.90).

The size of microglia also significantly increased in tri-cultures exposed to LPS as compared to vehicle control tri-cultures (Figure 2C, p = 0.0052). Increased microglia area is not consistent with *in vivo* observations, as activated microglia *in vivo* typically exhibit reduced area as they lose their ramifications and take on an amoeboid morphology (Fernández-Arjona et al., 2017). However, the morphology of the microglia following exposure to LPS is consistent with previous *in vitro* experiments, with the microglia showing a distinct polarity and the formation of large lamellipodia (Abd-El-Basset and Fedoroff, 1995; Nakamura et al., 1999; Persson et al., 2005). This flattening and formation of lamellipodia account for the increase in microglia area. We believe that this difference may not be due to a difference in microglia function but a result of the change from a 3D to 2D environment. In both instances, microglia become activated and prepare to migrate towards the site of injury. In the 3D environment, this involves the transformation from a ramified to amoeboid morphology to allow for the free movement through the parenchyma, while in the 2D environment this involves the extension of lamellipodia.

### Response to mechanical injury

In order to simulate a mechanical injury, we performed a scratch assay by drawing a pipette tip through co- and tri-cultures at DIV 7. Scratch assays are a common method used to model and measure cell migration (Liang et al., 2007), and have also been used to simulate mechanical injuries on cultured neurons and glial cells (Faber-Elman et al., 1996; Hirano et al., 2004; Tecoma et al., 1989). At 48 h following the scratch injury, we see a significant population of microglia migrated into the damaged area (Figure 3A). This is similar to what is observed *in vivo* where microglia are one of the first cells to respond to mechanical trauma in the CNS, and migrate towards the injury site to phagocytose cell debris and release pro-inflammatory cytokines (Corps et al., 2015; Davalos et al., 2005). In the tri-culture injury model, while the microglia that migrate into the injury site do not have a statistically significant larger surface area than the microglia in the control condition (Figure 3C, p = 0.078), they do show a trend of having a slightly larger surface area than microglia in the non-injured tri-culture. This increase in the surface area of the microglia is likely due to the extension of lamellipodia as the microglia migrate into the scratched area.

**Figure 3:**
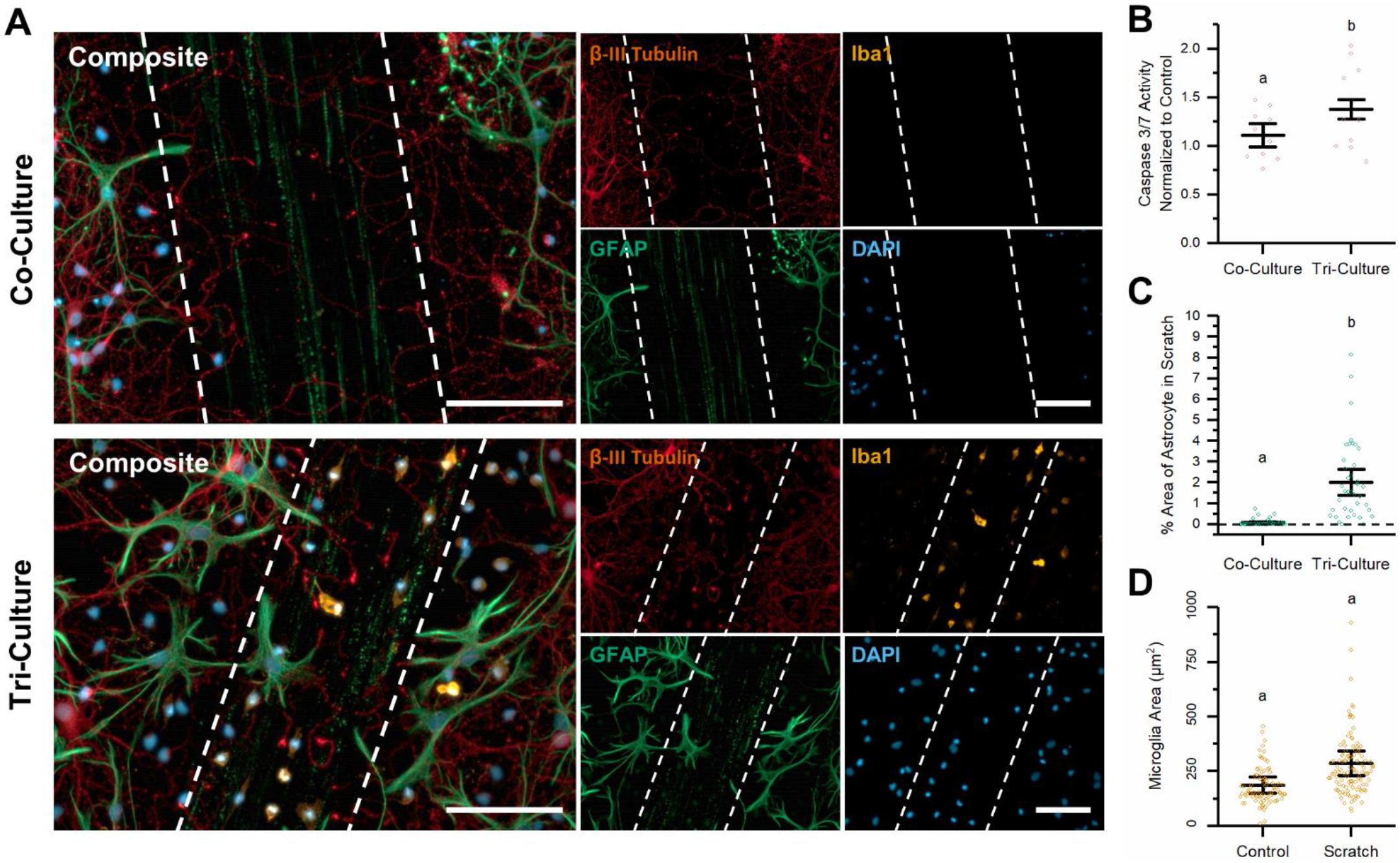
Response to mechanical injury. Comparing the effects from a simulated mechanical trauma (scratch) on tri- and co-cultures. (**A**) Representative fluorescence images of the tri- and co-cultures at 48 h following the simulated mechanical trauma. The dashed lines in the image highlight the area that was damaged by the scratch. The cultures were immunostained for the three cell types of interest: neurons–anti-βIII-tubulin (red), astrocytes–anti-GFAP (green), microglia–anti-Iba1 (orange) and the general nuclear stain DAPI (blue). Scale bar = 100 μM. (**B**) There was a significant increase in caspase 3/7 activity in the tri-culture as compared to the co-culture 48 h after the simulated mechanical trauma. **(C)** Comparing the percent area coverage of astrocytes in the scratched area between the co- and tri-cultures 48 h following the scratch.(**D**) Difference in the microglia area in the control and scratched tri-cultures. (**B-D**) All graphs display mean ± SD (n = 3), while the points indicate the values of the technical replicates. The letters above the bars indicate statistically significant differences (p < 0.05) as found by a t-test assuming unequal variances.

The neuroinflammatory response to traumatic brain injury (TBI) is a highly complex process, with both microglia and astrocytes responding to alarmins released by damaged cells to initiate an inflammatory cascade. The nature of this inflammatory response (neurotoxic or neuroprotective) is not well understood, and the details of the experimental or observational setup, including the time scale and type of injury, play a significant role in determining outcomes (Corps et al., 2015; Donat et al., 2017; Karve et al., 2016; Kawabori and Yenari, 2015). Furthermore, the exact role that microglia play in this process remains unknown. Activated microglia display a wide range of phenotypes with overlapping gene expression profiles and microglia polarization is no longer classified along a binary “M1 (neurotoxic) /M2 (neuroprotective)” scale based on the presence of specific markers (Donat et al., 2017; Li and Barres, 2017). However, it is generally hypothesized that microglia initially play a neuroprotective role through the clearing of cellular debris and alarmins, but transition to a more neurotoxic role as they begin to release pro-inflammatory cytokines (Donat et al., 2017; Kawabori and Yenari, 2015). This is supported by experiments showing that reducing the microglial population following induction of a hippocampal lesion improves functional outcomes and reduces synaptic loss. Conversely, reduction of the microglial population at the time of the insult increases neuronal cell loss (Rice et al., 2015). At 48 h following the scratch injury, caspase 3/7 activity was increased in both the tri- and co-cultures; however, the presence of microglia appeared to amplify a neurotoxic inflammatory response to the scratch, leading to a greater increase in caspase 3/7 activity (Figure 3B, p = 0.035). This is in line with the reported timeframe of the transition of microglia from a more neuroprotective to a more neurotoxic role in TBI (Simon et al., 2017).

Glial scarring is another aspect of the inflammatory response to mechanical injury, and it consists of the formation of a glial sheath consisting mostly of reactive astrocytes that encapsulate the damaged tissue, isolating it from healthy tissue (Polikov et al., 2005). While the formation of a glial scar may take weeks to fully mature, reactive astrocytes begin to migrate towards injury site in less than 24 h (Polikov et al., 2005). We observed this in the tri-culture as astrocytes began to migrate into the scratched area; in contrast, in the co-culture, the percent area coverage by astrocytes in the scratched area was almost non-existent (Figure 3C, p = 0.032). We hypothesize that the microglia that migrate into the injury site secrete factors that lead to increased astrocyte migration towards the injury. One such factor is matrix metalloprotease 9 (MMP-9), which is secreted by microglia in response to neuroinflammatory factors (Hu et al., 2014) and has been implicated in the migration of astrocytes and initial formation of the glial scar (Hsu et al., 2008). We observed that MMP-9 is secreted in tri-cultures, but not in co-cultures in response to LPS (Figure 5A).

### Response to excitotoxicity

At DIV 7, tri- and co-cultures were exposed to varying concentrations of glutamate for 1 h to simulate an excitotoxic event. Following glutamate exposure, both tri- and co-cultures were maintained for 48 h in tri-culture medium to eliminate the additional factors in the tri-culture medium as potentially confounding factors in the response. In particular, TGF-β (Boche et al., 2003) and IL-34 (Luo et al., 2013) have been shown to be neuroprotective during excitotoxicity. We observed no microglia in the co-culture media at the end of the experiment, further confirming that the co-culture medium is incapable of supporting even an insignificant microglia population.

Our results suggest that microglia in the tri-culture play a significant neuroprotective role during excitotoxic events, in line with recent *in vivo* (Kato et al., 2016; Larochelle et al., 2015; Szalay et al., 2016) and hippocampal slice culture experiments (Masuch et al., 2015; Vinet et al., 2012). We observed significant neuronal cell loss and astrocyte hypotrophy in neuron-astrocyte co-cultures treated with 25 μM glutamate, which is significantly reduced in similarly treated tri-cultures (Figure 4A). As glutamate concentrations increase, astrocytes in the co-culture became progressively more reactive, evidenced by increasing hypertrophy and loss of processes. This resulted in a significant increase in astrocyte surface area following treatment with 10 μM and 25 μM glutamate, which is not observed in the tri-culture (Figure 4B, p = 0.0022 and p = 0.0010 respectively). In order to quantify neuronal cell viability, we compared the percent area of the field-of-view stained for β-III tubulin with a circularity less than 0.2. The circularity cut-off was used to eliminate cell debris from apoptotic/necrotic neurons, which still stained for β-III tubulin, but lacked long cellular processes, and therefore had high circularity values. Across all concentrations of glutamate that were tested (5, 10 and 25 μM), there was significantly more neuronal cell loss in the co-culture then in the tri-culture (Figure 4C, p = 0.017, p = 0.0010 and p = 0.0017 respectively). Furthermore, we observed that the neuron percent area significantly decreased in the co-cultures upon exposure to increasing concentrations of glutamate. In contrast, significant decreases in the neuron percent area were observed in tri-culture only at the highest concentration of glutamate.

**Figure 4:**
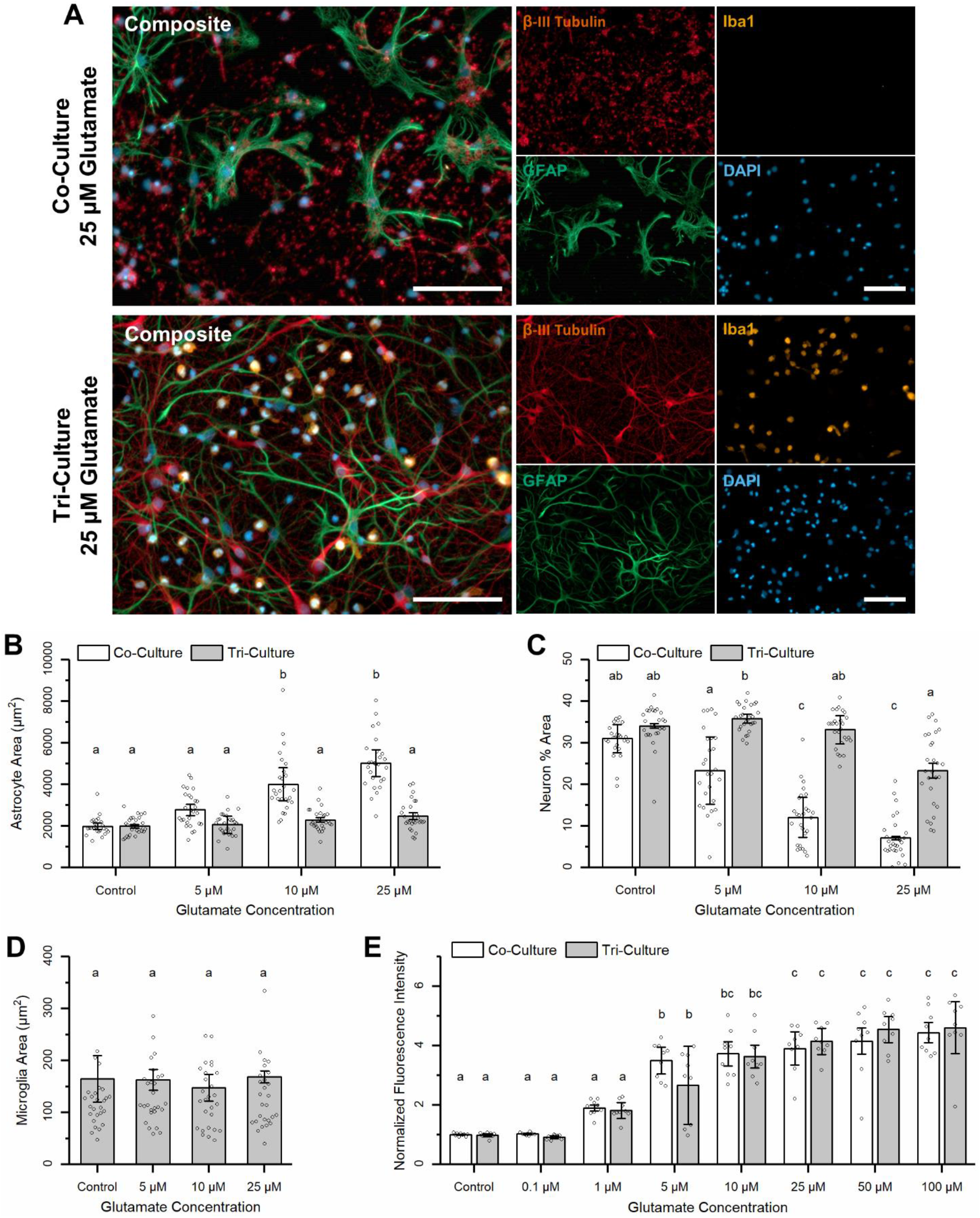
Response to excitotoxicity. (**A**) Representative fluorescence images of the tri- and co-cultures at 48 h following exposure to 25 μM glutamate for 1 h. The cultures were immunostained for the three cell types of interest: neurons–anti-βIII-tubulin (red), astrocytes–anti-GFAP (green), microglia–anti-Iba1 (orange) and the general nuclear stain DAPI (blue). Scale bar = 100 μM. (**B**) Comparing the average astrocyte area between the tri- and co-cultures following challenges with different concentrations of glutamate. A full analysis of the simple main effects can be found in supplementary data table 1. (**C**) Comparing the neuron percent area coverage, with a 0.2 circularity cut-off to eliminate cell debris, following excitotoxic challenge. A full analysis of the simple main effects can be found in supplementary data table 2. (**D**) The average microglia surface area did not change following treatment with different concentrations of glutamate. (**E**) Calcium imaging results showing the change in fluorescence intensity following treatment with different concentrations of glutamate. (**B-E**) All graphs display mean ± SD (n = 3), while the points indicate the values of the technical replicates. The letters above the bars indicate statistically significant differences (p < 0.05).

Surprisingly, unlike exposure to LPS or mechanical trauma, the exposure of the tri-cultures to glutamate did not appear to change the morphology of the microglia (Figure 4D). Specifically, the average microglia area did not change following treatment with different concentrations of glutamate (p = 0.81). However, this result is in line with previous research suggesting morphologically resting/ramified microglia can serve a neuroprotective role in *N*-methyl-D-aspartic acid (NMDA)-induced excitotoxicity in hippocampal slice cultures (Vinet et al., 2012).

We also confirmed that the tri-culture was electrophysiologically active via calcium imaging. For both the co- and tri-cultures, we observed spontaneous calcium influxes at DIV 7. Additionally, we compared the response of both culture types to varying concentrations of glutamate (Figure 4E). Analysis of the main effects did not identify a significant difference in the response of the tri-culture and co-cultures to different concentrations of glutamate (p = 0.78). Additionally, for both culture types, we observed a significant change in fluorescence intensity across a wide range of glutamate concentrations (p = 1.73 x 10^−15^) indicating that the neuroprotective effect was a result of the presence of microglia in the tri-culture, and was not due to a depressed neuronal response to glutamate.

### Tri-culture cytokine profile

We compared the cytokine secretion profiles of the control and LPS-exposed tri- and co-cultures using the proteome profiler rat XL cytokine array (Bio-Techne). Of the 79 cytokines detected by the array, 34 were detected at a relative concentration greater than 10% of the maximum in at least one of the samples. A hierarchical cluster analysis was performed on the results from these 34 cytokines (Figure 5A), which revealed three distinct expression profiles. In general, these profiles consist of cytokines secreted by both the co- and tri-cultures (green) and those only secreted by the tri-culture. This second group of cytokines can be further subdivided into cytokines that are secreted in relatively equal concentrations by the control and LPS-exposed tri-cultures (orange) and cytokines having increased expression in the LPS-exposed tri-cultures (purple). As expected, the co-cultures did not respond to LPS, leading to a nearly indistinguishable cytokine profile as compared to the control co-culture exposed only to vehicle.

**Figure 5:**
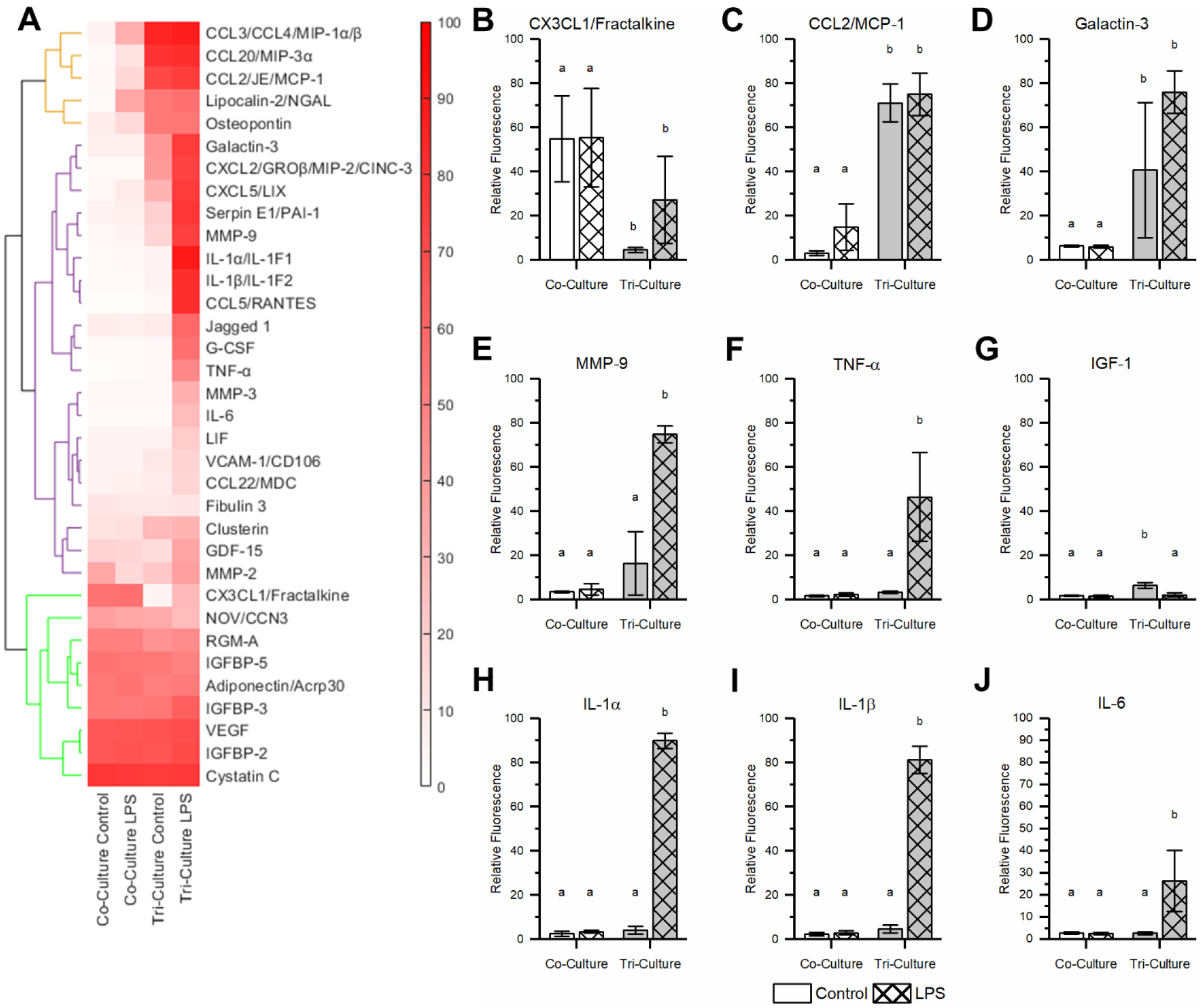
Tri-culture proteomic profile. Comparing the proteomic profile of the conditioned media from tri- and co-cultures after 48 h incubation with 5 μM LPS. (**A**) Heat map showing the relative cytokine concentrations scaled from 1 to 100. Proteins with a relative concentration of less than 10% of the maximum for all treatments were not included in the heat map, but can be found in (Figure 5, Supplement 1). Hierarchical cluster analysis revealed three major groups of cytokines consisting of cytokines present in the conditioned media from all culture types (green), cytokines with increased expression in both control and LPS challenged tri-culture (orange) and cytokines with increased expression only in the LPS exposed tri-cultures (purple). (**B-J**) The relative concentrations of specific cytokines of interest. All graphs display mean ± SD (n = 3). The letters above the bars indicate statistically significant differences (p < 0.05, 2-way ANOVA with the Tukey test used to analyze the simple main effects if necessary). A full breakdown of the statistical analysis can be found in supplementary data table 1.

All cytokines secreted by both the tri- and co-cultures have been shown to be expressed by either neurons or astrocytes in non-inflammatory conditions (Korecka et al., 2017; Lewitt and Boyd, 2019; Mathews and Levy, 2016; Rosenstein et al., 2010; Su et al., 2001), with the exception of adiponectin, which is found in the CSF of healthy individuals but has not been shown to be expressed by neurons or astrocytes (Yang et al., 2015). Additionally, with the exception of CX3CL1, these cytokines appear to be constitutively expressed by astrocytes and microglia as neither the media type nor the addition of LPS had an impact on their concentration in the conditioned media. CX3CL1 is expressed primarily by neurons in the CNS, and is found in both a membrane-bound and soluble form. Membrane bound CX3CL1 is important for microglia regulation, and thought to hold the microglia in resting state, while soluble CX3CL1 acts as a powerful chemoattractant for microglia and is also secreted by glial cells during neuroinflammatory conditions (Arnoux and Audinat, 2015; Paolicelli et al., 2014; Szepesi et al., 2018). Interestingly, we see a significant decrease in CX3CL1 concentration in the tri-culture as compared to the co-culture (Figure 5B, p = 0.0051). As microglia are an integral part of CNS homeostasis, the high concentration of CX3CL1 in the co-culture might be a compensatory response to the lack of microglia as co-cultures attempt to recruit them. The lack of CX3CL1 in the conditioned media from the control tri-cultures suggests that the majority of CX3CL1 in tri-cultures is membrane-bound and is interacting with the microglia in the tri-culture to hold them in a non-activated state. Additionally, we observe the presence of IGF-1 in the control tri-culture conditioned media that is not present in either the co-culture or the LPS-exposed tri-culture (p = 0.0010 and p = 0.0010 respectively). IGF-1 has been indicated as a trophic factor produced by microglia in postnatal (P3-7) mice and significantly increases the survival of layer V cortical neurons (Ueno et al., 2013). These results support previous conclusions that the overall health of the neurons in the tri-culture are not negatively impacted by the presence of microglia. Furthermore, we observe a significant decrease in caspase 3/7 activity in the tri-culture as compared to the co-culture (Figure 1 – figure supplement 2, p = 0.0031), which suggests that the health of neurons may be improved in the tri-culture. However, we are unable to determine the extent to which the presence of microglia versus the additional factors present in the tri-culture medium improve neural health.

Of the cytokines present in the tri-culture conditioned media that do not increase in expression in response to LPS, all have been shown to be secreted by microglia, and are also linked to neuroinflammatory states (Ambrosini et al., 2003; Goldmann and Prinz, 2013; Jha et al., 2015; Schindowski et al., 2011; Yu et al., 2017). In the case of CCL2, CCL3, and CCL20, the concentration of these cytokines in the conditioned media are at the upper limit of the array under control conditions, thus, an increase in concentration in response to LPS may not have been detected. However, in the case of lipocalin-2 and osteopontin, we would expect their concentrations to increase following LPS insult. The fact that this was not observed may indicate that the tri-culture is in a slightly inflamed state, possibly due to trauma associated with dissection to isolated cells from the intact brain. However, there are a significant number of pro-inflammatory cytokines that are present in the conditioned media only after treatment with LPS, including many of the hallmark pro-inflammatory cytokines secreted by activated microglia. In particular, levels of TNF-α, IL-1α, IL-1β and IL-6 are significantly increased in the conditioned media of tri-cultures challenged with LPS (Figure 5F-J, p = 0.0033, p = 0.0010, p = 0.0010, p = 0.013 respectively). These four cytokines are consistently secreted by microglia in response to LPS in a variety of experimental conditions and are often used as biomarkers to indicate neuroinflammatory or neurodegenerative disorders (Kothur et al., 2016). Furthermore, the increase in the concentration of CX3CL1 coupled with the decrease in IGF-1 concentration in LPS-exposed tri-culture conditioned media further supports conclusions from the studies described above indicating that the microglia in the tri-culture are responding to LPS in a pro-inflammatory manner. Ultimately, these results support the use of this tri-culture as a neuroinflammatory model, as exposure to LPS leads to the secretion of a significant number of pro-inflammatory cytokines associated with microglia and responses to LPS treatments that are not present in the co-culture condition.

## Conclusions

Here, we describe an *in vitro* model consisting of a tri-culture of neurons, astrocytes and microglia, that more faithfully mimics neuroinflammatory responses than standard mono- and co-cultures. This tri-culture is established by culturing primary cortical cells from neonatal rats in a tri-culture medium supplemented with 100 ng/mL IL-34, 2 ng/mL TGF-β and 1.5 μg/mL cholesterol. We characterized the tri-culture under a number of neuroinflammatory scenarios, including exposure to LPS, mechanical injury and excitotoxic challenge. We compared the response of the tri-cultures to that of the neuron-astrocyte co-cultures to these different neuroinflammatory insults and found that the presence of microglia changes the response in each case. Moreover, the response of the tri-culture to these neuroinflammatory challenges are more in line with what is reported *in vivo* and in slice culture models.

Microglia in the tri-culture model displayed an amoeboid morphology and not the classical ramified morphology of phenotypically “resting” microglia in the adult CNS. We believe this is due to the fact that the primary microglia present in the culture are derived from neonatal rats, and therefore are still displaying their neonatal morphology and phenotype (Cuadros and Navascue, 1998; Dalmau et al., 2003; Harry and Kraft, 2012). This hypothesis is supported by the presence of galactin-3 in the tri-culture, which has been implicated as one of the factors that maintains the amoeboid morphology of microglia in early postnatal rats (Reichert and Rotshenker, 2019), along with the presence of IGF-1, which is produced by microglia in early postnatal mice (Ueno et al., 2013). However, the proteome profile of the tri-culture revealed the presence of cytokines typically associated with activated microglia, indicating that the tri-culture may be in a somewhat inflamed state. We have demonstrated that microglia in the tri-culture can assume either neurotoxic and neuroprotective phenotypes depending on the inflammatory challenge, and these phenotypes are consistent with *in vivo* observations (Kato et al., 2016; Polikov et al., 2005). In particular, the tri-culture shows a classic neuroinflammatory response to LPS, with the secretion of the pro-inflammatory cytokines TNF-α, IL-1β and IL-6. In addition, we observed that in tri-cultures challenged with LPS, astrocytes became hypertrophic coincident with an increase in caspase 3/7 activity. Conversely, in response to an excitotoxic challenge, the microglia in the tri-culture are neuroprotective, reducing neuronal cell death following glutamate-induced excitotoxicity.

We believe this tri-culture has multiple advantages over other methods used to study neuroinflammation *in vitro*. The most obvious benefit is the presence of neurons, astrocytes and microglia in the same cell culture, which will allow researchers to study the complex interplay between these cells that leads to different responses to inflammatory stimuli. Additionally, the use of a single culture (as opposed to using conditioned media to mimic the influence of cell types absent in the culture) allows for the observation of cell-cell interactions and other mechanisms with spatiotemporal nuances that may be lost when using models involving multiple different types of cultures. Another major benefit of the tri-culture model is its relative simplicity. The only modification needed to establish and maintain the tri-culture model is the use of a specialized tri-culture medium. Compared to other models that involve multiple different culture models or the addition of cells at specific time points during culture, the simplicity of this tri-culture model lends itself to high-throughput experiments, and, therefore, it may be an effective tool in the early screening of potential therapeutic molecules. Additionally, this tri-culture model is amenable to experiments involving more complex culture setups, such as microfluidic and organ-on-a-chip devices. Ultimately, we believe that the neuron, astrocyte and microglia tri-culture described in this paper can be a useful tool to study neuroinflammation *in vitro* with improved accuracy in predicting *in vivo* neuroinflammatory phenomena.

## Experimental

### Culture Media Preparation

Base media (plating medium and co-culture medium) were prepared as previously described (Chapman et al., 2015). Briefly, plating medium consisted of Neurobasal A culture medium supplemented with 2% B27 supplement, 1x Glutamax, 10% heat-inactivated horse serum and 1 M HEPES at pH 7.5, while the co-culture medium consisted of Neurobasal A culture medium supplemented with 2% B27 supplement and 1x Glutamax (all from ThermoFisher). The tri-culture medium consisted of supplementing the co-culture medium with 100 ng/mL mouse IL-34 (R&D Systems), 2 ng/mL TGF-β (Peprotech) and 1.5 μg/mL ovine wool cholesterol (Avanti Polar Lipids). Due to the limited shelf life of the IL-24 and TGF-β, tri-culture medium was made new each week.

### General cell culture

All procedures involving animals were conducted in accordance with the NIH Guide for the Care and Use of Laboratory Animals following protocols approved by the University of California, Davis Institutional Animal Care and Use Committee. Timed-pregnant Sprague Dawley rats were purchased from Charles River Laboratory (Hollister, CA). All animals were housed in clear plastic shoebox cages containing corn cob bedding under constant temperature (22 ± 2 °C) and a 12 h light-dark cycle. Food and water were provided ad libitum. Primary cortical cell cultures were prepared from postnatal day 0 rat pups as previously described (Wayman et al., 2012). Neocortices from all pups in the litter were pooled, dissociated and plated at a density of 650 cells/mm^2^ on substrates precoated with 0.5 mg/mL of poly-L-lysine (Sigma) in B buffer (3.1 mg/mL boric acid and 4.75 mg/mL borax, Sigma) for 4 h at 37 °C and 5% CO_2_ then washed with sterile deionized water and covered with plating medium. Primary cortical cells were plated in plating medium, and allowed to adhere for 4 h before the medium was changed to the co- or tri-culture medium. Half-media changes were performed at DIV 3, 7 and 10 with the respective media types.

### Neuroinflammatory challenges

To simulate bacterial infection, cultures were challenged with LPS (3.0 x 10^6^ EU/mg; Sigma) was reconstituted in sterile Dulbecco’s Phosphate Buffered Saline solution (DPBS) with calcium and magnesium (DPBS+) (Sigma) to a stock solution of 1 mg/mL and stored at −20°C. Following the DIV 7 media change, each well was spiked with LPS solution to a final concentration of 5 μg/mL or an equal volume of sterile DPBS+ to act as the vehicle control. To simulate mechanical injury, a scratch was made in the tri- and co-cultures following the DIV 7 media change. A cross (~200-300 μm wide) was scratched in the center of each well using a sterile 200 μL micropipette tip. Excitotoxicity was triggered by adding varying concentrations of glutamate to the cultures. Prior to each experiment, a fresh 50 mM solution of L-glutamic acid (Sigma) in DPBS+ was prepared. This 50 mM solution of L-glutamic acid was diluted with sterile DPBS+ to 100X stocks. At DIV 7, half of the medium was removed from each well and stored at 37°C. The glutamate solutions were diluted 1:100 directly into the cultures; vehicle controls received an equal volume of sterile DPBS+. Glutamate-exposed cultures were incubated at 37 °C for 1h. For co-cultures, the media collected from the cultures prior to the addition of glutamate was combined with an equal volume of tri-culture medium that had twice the concentration of supplemental factors (4 ng/mL TGF-β, 200 ng/mL IL-34 and 3 μg/mL cholesterol) to create a tri-culture medium that contained any secreted factors from the co-culture, while the stored tri-culture media was combined with an equal volume of standard tri-culture medium. Following the 1 h incubation, the medium from each well was completely removed and quickly replaced with the appropriate medium type for the culture condition.

### Immunostaining

At the conclusion of an experiment, cell cultures were washed 3 times with 37°C DPBS+, and were fixed using 4% w/v paraformaldehyde in PBS (PFA; Affymetrix) for 1 h. The fixed cells were first washed twice with 0.05% v/v Tween20 (Sigma) solution in DPBS+, followed by a 3 min permeabilization with 0.1% v/v Triton X-100 (ThermoFisher) solution in DPBS+ and two additional washes with the Tween20 solution. Samples were blocked with a solution of 5% v/v heat-inactivated goat serum (ThermoFisher) and 0.3 M glycine (Sigma) in DPBS+ (blocking buffer) for 1 h. Following this blocking step, samples were incubated for 1 h in primary antibody solution containing mouse anti-β-III tubulin (1:500 dilution, ThermoFisher), rabbit anti-GFAP (1:100 dilution, ThermoFisher) and chicken anti-Iba1 (1:500 dilution, Abcam) in blocking buffer. Samples were then washed 3 times with Tween20 solution before a 1 h incubation with secondary antibody solution containing goat anti-mouse conjugated to AlexaFluor 647 (1:500 dilution, ThermoFisher), goat anti-rabbit conjugated to AlexaFluor 488 (1:500 dilution, ThermoFisher) and goat anti-chicken conjugated to AlexaFluor 555 (1:500 dilution, ThermoFisher) in DPBS+. Following incubation with secondary antibody solution, the samples were washed 3 times with DPBS+. Lastly, samples were incubated for 5 min with a 4’,6-diamidino-2-phenylindole (DAPI) solution (1:20000 dilution in DI H_2_O, Sigma), followed by an additional Tween20 solution wash before mounting onto glass slides using ProLong Gold Antifade Mountant (ThermoFisher).

### Morphological Analysis

For morphological analysis, cultures were fixed with a 4% w/v paraformaldehyde solution in PBS and immunostained as described above. All samples were acquired with a Zeiss Observer D1 inverted fluorescent microscope at 100x or 200x magnification and analyzed using ImageJ. The cell number/mm^2^ of the different cell types was determined by manually counting the number of nuclei that were co-localized with β-III tubulin (neurons), GFAP (astrocytes) or Iba1 (microglia) from the 100x magnification images. The average astrocyte/microglia areas were determined by manually tracing the outline of astrocytes/microglia from 200x magnification images and determining the area inside the trace. For both of these manual quantification methods, the images were de-identified and the investigator was blinded to the experimental group. Percent area coverage of neurons or astrocytes was determined through the use of the Huang auto-thresholding method (Huang and Wang, 1995) on the β-III tubulin (neurons) or GFAP (astrocytes) channel.

### Apoptosis

Apoptosis was quantified using the Caspase-Glo^®^ 3/7 Assay System (Promega) according to the manufacturer’s protocol. Luminescence was measured using a H1 hybrid microplate reader (BioTek Instruments).

### Calcium Imaging

Prior to imaging, tri- and co-cultures were loaded with cell-permeant Fluo-4 AM calcium indicator (ThermoFisher) following the manufacturer’s protocol. To determine the effect of glutamate on calcium fluxes in the tri- and co-cultures, each culture type was spiked with varying concentrations of L-glutamic acid (Sigma) in DPBS+ or an equal volume of DPBS+. For each well, prior to the addition of the glutamate solution, a 200x magnification fluorescent image was taken, after which the shutter to the light source was closed and the glutamate solution was added to the well. Following a 2 min incubation in the glutamate solution, a second fluorescent image was taken over the same field-of-view with the same exposure time, and was used to compare the change in fluorescence intensity following the addition of glutamate.

### Proteome Profile

Following the DIV 7 media change, co- or tri-cultures were incubated with 5 μg/mL LPS or vehicle control for 48 h. Following incubation, the conditioned media was spun down to remove any cells, and the supernatant containing the conditioned media was stored at −80 °C until analyzed. The proteome profile was determined using a Proteome Profiler Rat XL Cytokine Array (Bio-Techne) in conjugation with the IRDye^®^ 800CW (Bio-Techne) for use with the LI-COR Odyssey® Imaging System. Relative concentrations of each cytokine in conditioned media was determined using ImageJ to compare the total pixel intensity from each spot. Hierarchical cluster analysis was performed using the MATLAB (2019a) bioinformatics toolbox.

### Statistical Methods

For all experiments, a minimum of three biological replicates was used with a minimum of three technical replicates per biological replicate. Furthermore, unless otherwise noted, for experiments requiring image analysis at least three predetermined fields were analyzed per technical replicate to account for variability within the culture itself. When comparing the response of the tri- and co-cultures to different treatments, a 2-way ANOVA was used. If the interaction was determined not significant (p < 0.05), then the analysis of the main effects were used to compare the two treatments. If a significant interaction was found, analysis of the simple main effects was conducted via a *post hoc* Tukey test. A one-way ANOVA test was used when comparing multiple groups against a single treatment, while a two-tailed Student’s t-test assuming unequal variances was used when only two groups were analyzed. For all experiments, statistical significance was determined by p-values < 0.05.

## Acknowledgements

We would like to thank Sunjay Sethi, Harmanpreet Panesar, and Felipe Da Costa Souza (University of California, Davis) for their help in obtaining primary cortical cells, along with Barath Palanisamy for his help in image acquisition and de-identification. We acknowledge funding from the National Science Foundation (CBET&DMR-1454426), the National Institutes of Health (R21 EB024635), and University of California - Microbiome Special Research Program. N. Goshi was supported by the UC Davis Biotechnology Training Program award.

