## Supplemental Information for "A Primary Neural Cell Culture Model for Neuroinflammation"

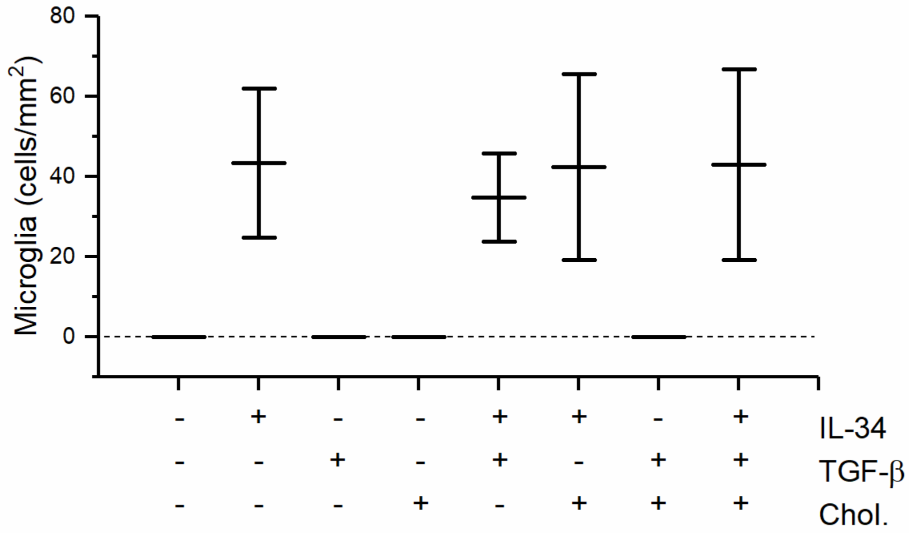


**Figure 1, Supplementary Figure 1:** Tri-culture media supplement requirements for microglia survival at DIV 7. The results indicate that IL-34 is required for microglial survival in the tri-culture. The figure shows the mean ± SD of the technical replicates (n = 4) of a single biological replicate.

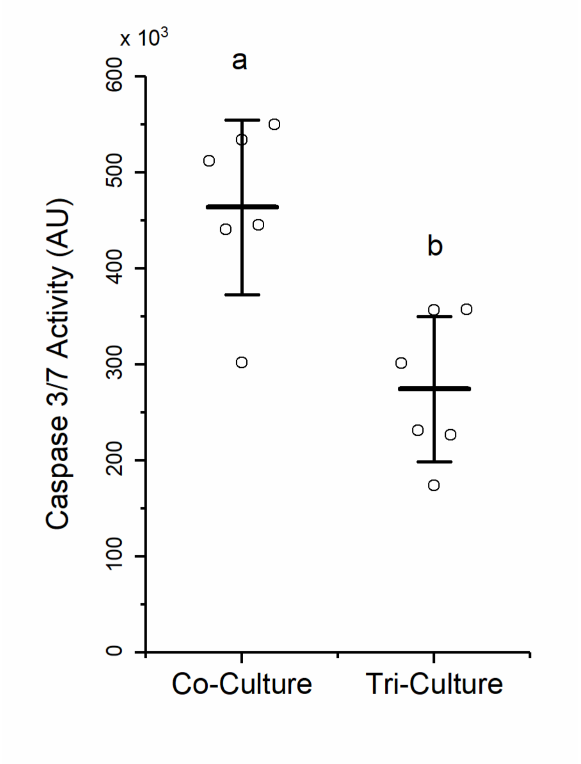


**Figure 1, Supplementary Figure 2:** The tri-culture shows reduced caspase 3/7 activity at DIV 9 ($n = 6$). The letters above the bars indicate statistically distinct groups ($p < 0.05$), while the points indicate the values of the technical replicates.

| | | Co-Culture | | | | Tri-Culture | | | |
|---|---|---|---|---|---|---|---|---|---|
| | | Control | 5 µM | 10 µM | 25 µM | Control | 5 µM | 10 µM | 25 µM |
| Co-Culture | Control | | | | | | | | |
| | 5 µM | 0.31 | | | | | | | |
| | 10 µM | 0.0010 | 0.036 | | | | | | |
| | 25 µM | 0.0010 | 0.0010 | 0.11 | | | | | |
| Tri-Culture | Control | 0.90 | 0.33 | 0.0010 | 0.0010 | | | | |
| | 5 µM | 0.90 | 0.44 | 0.0010 | 0.0010 | 0.90 | | | |
| | 10 µM | 0.90 | 0.81 | 0.0022 | 0.0010 | 0.90 | 0.90 | | |
| | 25 µM | 0.78 | 0.89 | 0.0063 | 0.0010 | 0.81 | 0.90 | 0.90 | |

**Figure 4, Supplementary Data Table 1:** Analysis of the simple main effects (Tukey test) from Figure 4B. The p-values from each pairwise result are shown on the table, with p-values less than 0.05 highlighted in green.

| | | Co-Culture | | | | Tri-Culture | | | |
|---|---|---|---|---|---|---|---|---|---|
| | | Control | 5 µM | 10 µM | 25 µM | Control | 5 µM | 10 µM | 25 µM |
| Co-Culture | Control | | | | | | | | |
| | 5 µM | 0.27 | | | | | | | |
| | 10 µM | 0.0010 | 0.038 | | | | | | |
| | 25 µM | 0.0010 | 0.0017 | 0.72 | | | | | |
| Tri-Culture | Control | 0.90 | 0.051 | 0.0010 | 0.0010 | | | | |
| | 5 µM | 0.74 | 0.017 | 0.0010 | 0.0010 | 0.90 | | | |
| | 10 µM | 0.90 | 0.087 | 0.0010 | 0.0010 | 0.90 | 0.90 | | |
| | 25 µM | 0.27 | 0.90 | 0.037 | 0.0017 | 0.051 | 0.017 | 0.087 | |

**Figure 4, Supplementary Data Table 2:** Analysis of the simple main effects (Tukey test) from Figure 4C. The p-values from each pairwise result are shown on the table, with p-values less than 0.05 highlighted in green.

| | Control | 0.1 µM | 1 µM | 5 µM | 10 µM | 25 µM | 50 µM | 100 µM |
|---|---|---|---|---|---|---|---|---|
| **Control** | | | | | | | | |
| **0.1 µM** | 0.9 | | | | | | | |
| **1 µM** | 0.68 | 0.56 | | | | | | |
| **5 µM** | 0.0010 | 0.0010 | 0.0023 | | | | | |
| **10 µM** | 0.0010 | 0.0010 | 0.0010 | 0.45 | | | | |
| **25 µM** | 0.0010 | 0.0010 | 0.0010 | 0.0380 | 0.90 | | | |
| **50 µM** | 0.0010 | 0.0010 | 0.0010 | 0.0016 | 0.29 | 0.90 | | |
| **100 µM** | 0.0010 | 0.0010 | 0.0010 | 0.001 | 0.087 | 0.65 | 0.90 | |

**Figure 4, Supplementary Data Table 2:** Analysis of the simple main effects (Tukey test) from Figure 4C. The p-values from each pairwise result are shown on the table, with p-values less than 0.05 highlighted in green.

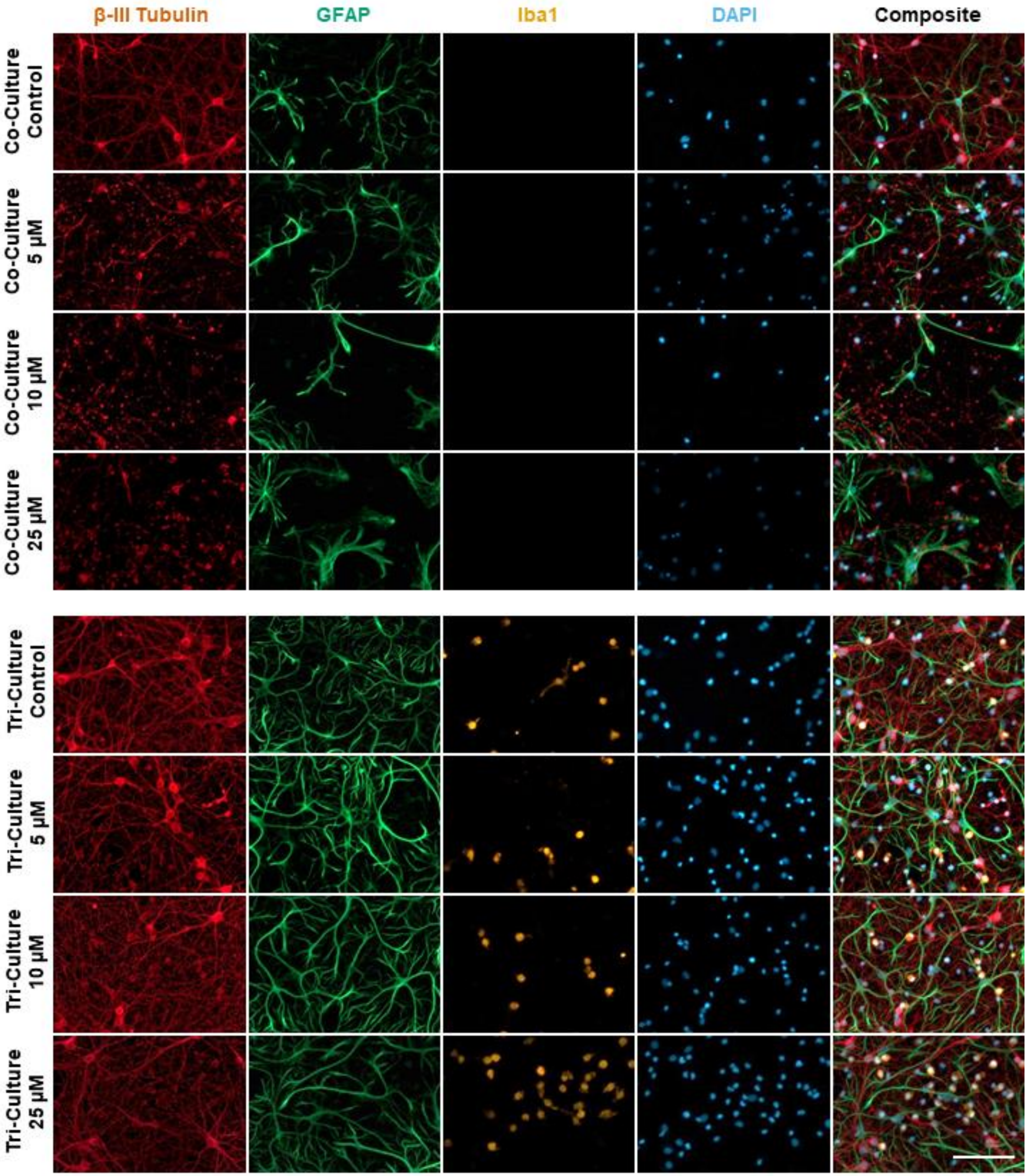


**Figure 4, Supplementary Figure 1:** Representative images of the tri- and co-cultures 48 h following a 1 h treatment with different concentrations of glutamate or vehicle control. The cultures were immunostained for the three cell types of interest: neurons – anti-βIII-tubulin (red), astrocytes – anti-GFAP (green), microglia – anti-Iba1 (orange) and the general nuclear stain DAPI (blue). Scale bar = 100 µM.

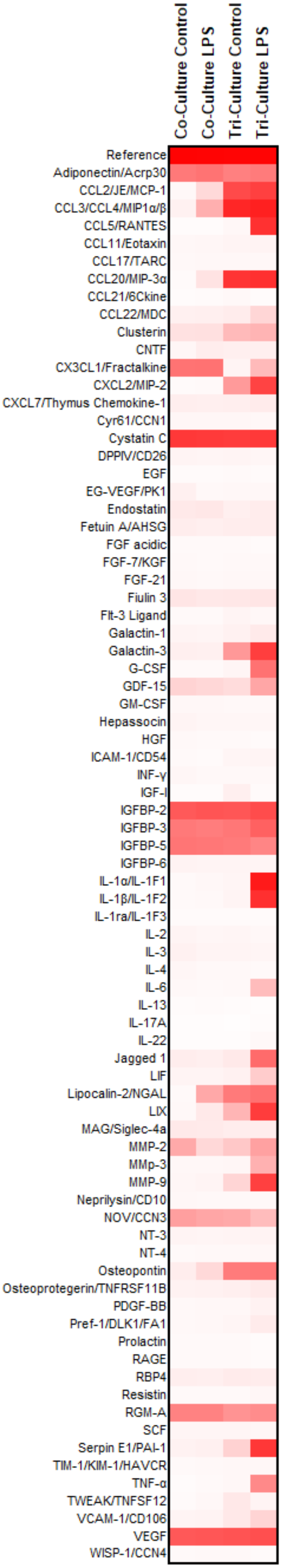


**Figure 5, Supplementary Figure 1:** Complete proteomic profile from Figure 5A.

| | | Main Effects | | Simple Main Effects | | | |
|---|---|---|---|---|---|---|---|
| | **Interaction** | **Co-Culture vs Tri-Culture** | **Control vs LPS** | **Co-Culture Control vs Tri-Culture Control** | **Co-Culture Control vs Co-Culture LPS** | **Tri-Culture Control vs Tri-Culture LPS** | **Co-Culture LPS vs Tri-Culture LPS** |
| **CXC3L1** | 0.31 | 0.0051 | 0.29 | N/A | N/A | N/A | N/A |
| **CCL2** | 0.43 | 0.00 | 0.14 | N/A | N/A | N/A | N/A |
| **Galactain 3** | 0.090 | 0.00050 | 0.095 | N/A | N/A | N/A | N/A |
| **MMP9** | 0.0020 | N/A | N/A | 0.23 | 0.90 | 0.0010 | 0.0010 |
| **TNF-α** | 0.0065 | N/A | N/A | 0.90 | 0.90 | 0.0033 | 0.0030 |
| **IGF-1** | 0.0024 | N/A | N/A | 0.0010 | 0.90 | 0.0010 | 0.76 |
| **IL-1α** | 4.46E-10 | N/A | N/A | 0.77 | 0.90 | 0.0010 | 0.0010 |
| **IL-1β** | 3.97E-08 | N/A | N/A | 0.82 | 0.90 | 0.0010 | 0.0010 |
| **IL-6** | 0.017 | N/A | N/A | 0.90 | 0.90 | 0.013 | 0.012 |

**Figure 5, Supplementary Data Table 1:** Statistical analysis of Figure 5B-J. The p-values from the 2-way ANOVA and simple main effects analysis (Tukey Test) are shown. p-values < 0.05 are highlighted in green.
